# WaveLimit: An optimal spike sorting inclusion boundary

**DOI:** 10.1101/2025.07.01.662629

**Authors:** Xavier Scherschligt, Wei-Hsien Lee, Colin A. Dallimore, Evan Schrader, Zheng Liu, Adam G. Rouse

## Abstract

Spike sorting is the process of assigning neuronal action potentials to individual putative neurons based on extracellular recordings. Spike sorting may be partitioned into five major components: i) raw neural data is filtered, ii) spiking events are extracted as waveforms, iii) features are extracted from the waveforms, iv) clusters of waveforms are defined, and v) individual waveforms are assigned to their respective clusters. Here, we focus on the often underappreciated fifth component, deriving a useful principle to define a cluster boundary to maximize the theoretical information available from a single neuron. We implemented this boundary, along with an automatic cluster identifier, in a novel spike sorting algorithm, WaveLimit. We then compared WaveLimit to three state-of-the-art spike sorters. WaveLimit identified either the same or more units and included more spiking events per unit than the other sorters. WaveLimit also found units with fewer inter-spike interval violations and higher signal-to-noise ratios. Thus, better defining the cluster boundary improved spike sorting.

## Main

Spike sorting is the process of putatively assigning waveforms to a particular neuron based on extracellular electrical recordings of neurons’ action potentials^1^. Each neuron generates a characteristic waveform every time it fires an action potential, which distinguishes them from waveforms generated by other neurons recorded from the same channel^2^. While some neurons’ waveforms are so large compared to background noise and the waveforms of other neurons that sorting is unambiguous, many are not. In ambiguous cases, waveforms are clustered and assigned to individual units most likely to represent each neuron. The advent of chronic recording arrays and multi-channel recording systems, however, has dramatically increased the number of channels encountered with less precise single unit isolation^3^. Historically, manual spike sorting was the gold standard, but it is impractical with many channels and is biased^4^.

The typical approach to automated spike sorting may be partitioned into five major components: i) the raw neural recording is bandpass filtered, ii) neuronal spiking events are extracted by applying an amplitude threshold, iii) relevant features are extracted from the waveforms of those spiking events, iv) automatic discovery of clusters is performed in the feature space, and v) individual waveforms are assigned to their respective clusters or discarded as noise (for a more in depth discussion, see ^1,5^). While the first four components have received significant attention, surprisingly little focus has been placed on the final sorting component compared to feature extraction and cluster discovery. Anecdotally, automatic spike sorters often appear to struggle from channel to channel by being either too inclusive or too restrictive in the number of waveforms added to a cluster. With most algorithms, the assignment boundary is either inherited from or closely tied to clustering, with parameters that are fixed across all units and channels^1,4,5^. The novel innovation of this present work is deriving a useful principle for determining how tightly or expansively to define a cluster boundary for each individual sorted unit.

Several methods have been identified for assessing the quality of a unit’s spike sorting and the likelihood all the sorted waveforms are action potentials from a single neuron. The distribution of observed waveforms can be estimated as the noise around the mean unit waveform and can be well-fit with a multivariate Gaussian^6^. The Gaussian fit provides an estimate of the fraction of spikes occurring within any given boundary. Furthermore, Gaussian fits for multiple units on the same channel allow for estimating the fraction of overlap. Additionally, any given neuron cannot fire two action potentials in quick succession. The biophysics underlying action potential generation create an absolute refractory period during which no second action potential can occur, followed by a relative refractory period during which a second action potential is less likely^7^. If the observed distribution of inter-spike intervals contains intervals within this refractory period (caused by rogue spikes from another unit or noise), the fraction of spikes not originating from the unit of interest can be estimated^8^.

These estimates of spike sorting quality provide useful metrics of sorting performance, but can they be leveraged to improve automated sorting algorithms themselves? Theoretically, one would like to include as many spikes from the source neuron as possible. However, it is often impossible to include all spikes from a single neuron without including some rogue spikes that may result from the superposition of noise and waveforms from other neurons. One is thus faced with a tradeoff between including as many true spikes as possible with a minimal number of rogue spikes.

Here, we present a theoretical framework for choosing a boundary and implement a new sorting algorithm focused on estimating this optimal boundary. This sorting principle for maximizing estimated true spikes from the isolated neuron while minimizing false spikes is derived assuming multivariate Gaussian distributions of clustered waveforms and Poisson process statistics of inter-spike intervals. We then describe WaveLimit^9^, our MATLAB implemented spike sorter based on these principles. Finally, we compare the performance of WaveLimit to that of three other prominent sorting algorithms.

## Results

### Sample channel sorting

Figure 1 shows an intracortical microelectrode recording from a single example channel. Figure 1a displays spike waveforms captured with a single negative threshold crossing. The waveforms were decomposed into their top two principal components which are projected in Figure 1b. In this principal component analysis (PCA) space, one cluster of waveforms stands out clearly, whereas other clusters overlap, suggesting the presence of additional units that are less distinct and noisier. WaveLimit initially identified three cluster centroids, a clearly identifiable cluster (top right of Fig 1a) along with two other clusters. The centroid positions are updated iteratively to achieve their final location.

**Figure 1.**
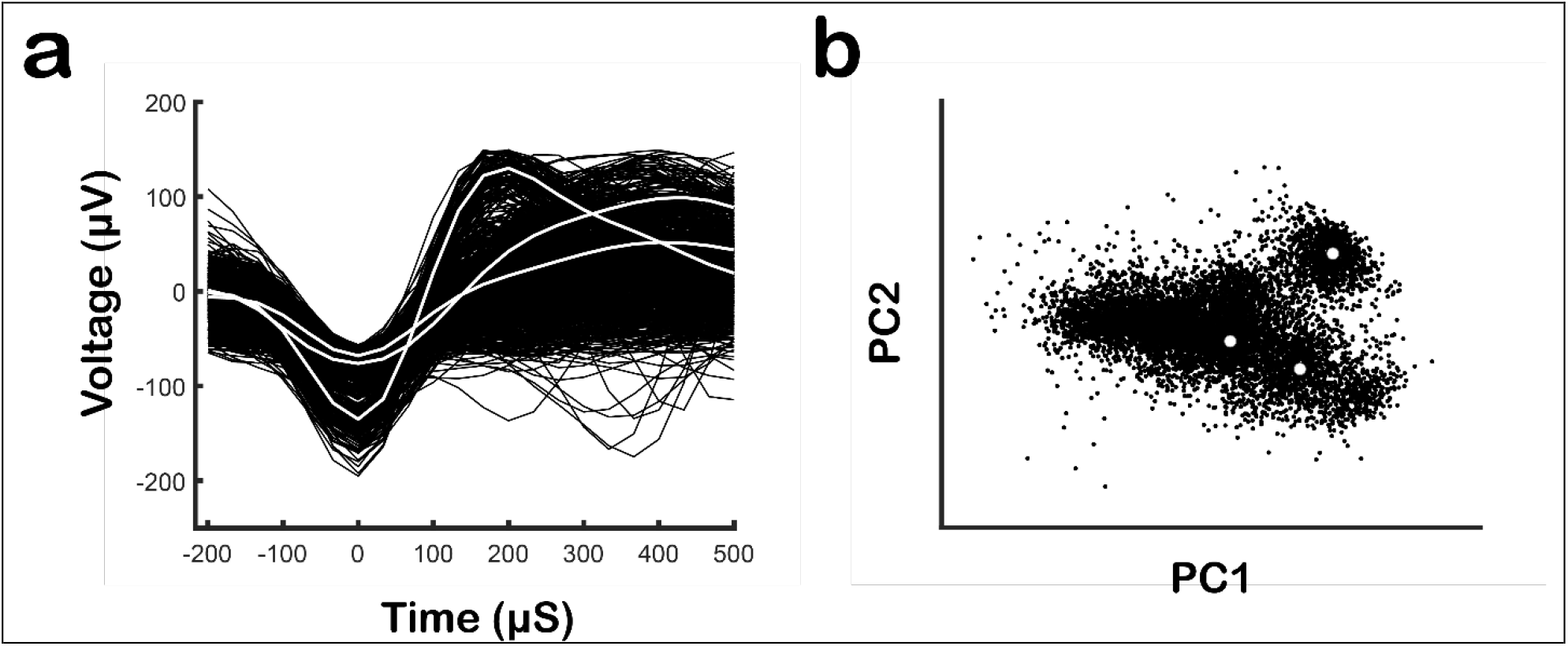
Spike Threshold and Waveform Feature Extraction. **(a)** Intracortical microelectrode recording for an example channel. The channel was bandpass filtered and only those electrical deviations which exceeded an amplitude threshold are shown. The resulting waveforms were aligned to the threshold crossing. **(b)** Top two principal components of the waveforms. The initial waveform cluster centroids identified by WaveLimit are shown in white. The average waveforms of those centroids are shown in white in Figure 1a.

Once centroids that define a single unit are identified, the waveforms can be added to the single unit based on their distance from the centroid starting with the closest. As one moves to waveforms farther away, the sorter must pick a limit to stop adding spikes. While we cannot confidently decide if any individual waveform is from the single neuron as we get farther from the centroid, we can estimate the probability that it is from the single neuron. This probability is decided by two factors: i) how many of the waveforms from the total number of action potentials generated by the neuron of interest were included and ii) what fraction of the waveforms are rogue waveforms from other neurons or noise.

We first estimate the number of waveforms arising from a single neuron using the Euclidean distance from each cluster centroid. Figure 2a’s upper panel plots the rank-ordered distances between each cluster centroid and the individual waveforms. We then fit a chi-squared (*χ*^2^) distribution to these observed distances, which assumes a Gaussian noise distribution of waveforms around the center waveform. This *χ*^2^ fit is performed on only the closest distances as almost all these waveforms likely originated from the single neuron defined by the centroid. As we move to distances farther from the cluster centroid, we can analyze when the distances and *χ*^2^ fit start to diverge, due to encountering waveforms from other neurons or noise. Using the *χ*^2^ fit, the cumulative fraction of waveforms included with each additional waveform distance can be estimated, as shown in Figure 2a’s lower panel.

**Figure 2.**
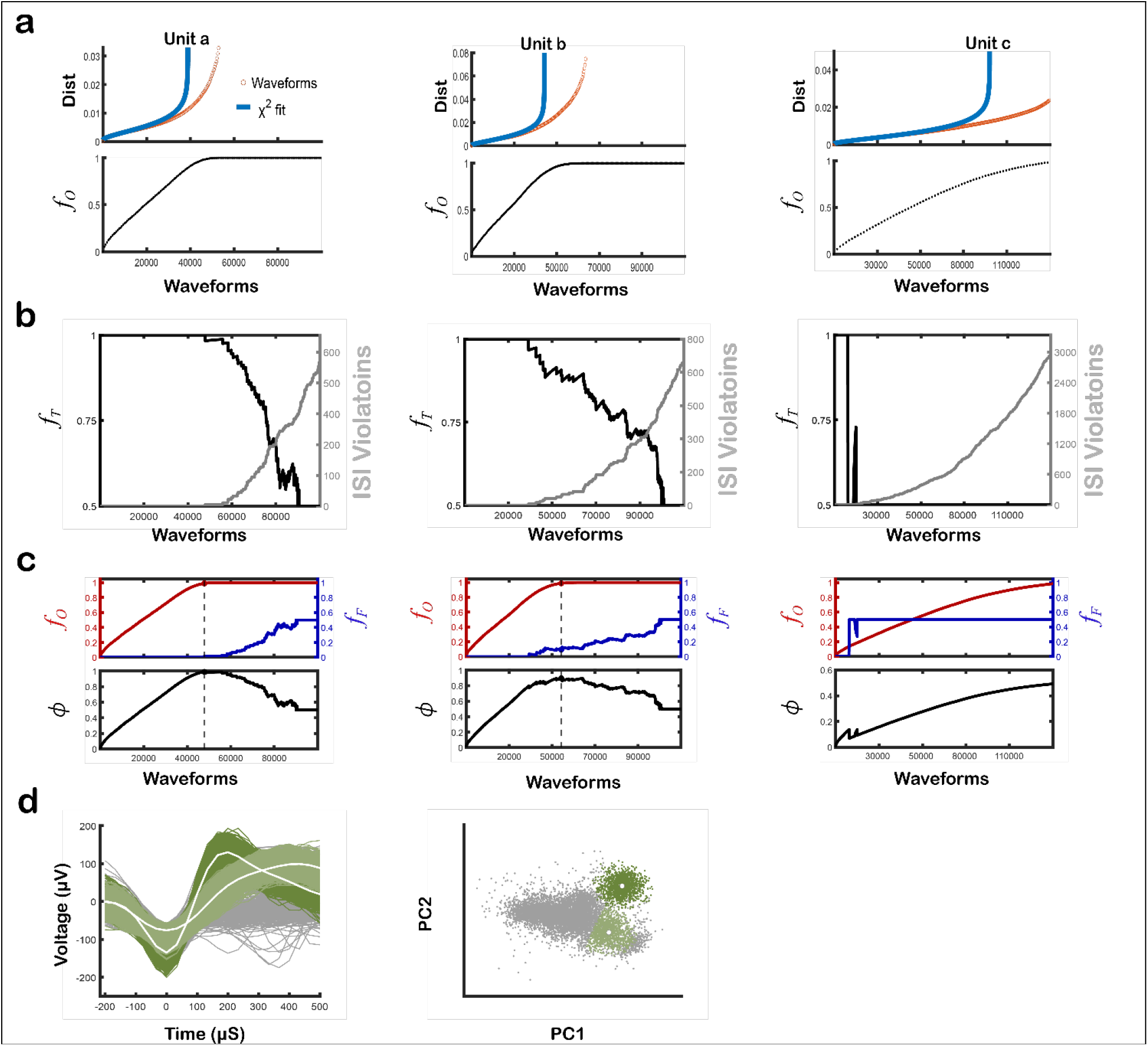
Optimal Spike Inclusion Boundary. **(a)** The top panel shows the Euclidean distance between each spike and the spike cluster centroid, sorted from smallest to largest (orange), for Units a, b, and c. The blue line represents the *χ*^2^fit to that distribution for both units. The lower panel illustrates the cumulative density function (called the fraction of observed waveforms: *f*_O_) of the *χ*^2^ fit for each unit. **(b)** The number of inter-spike interval violations incurred is shown in gray, measured at increasing radial bin distances from each unit’s centroid. The fraction of true spikes (*f*_T_) from these ISI violations is shown in black. **(c)** The fraction of observed spikes (*f*_O_) is shown in red, and the fraction of false spikes (*f*_F_) is shown in blue, measured at increasing radial bin distances from each unit centroid. The optimized spike inclusion boundary is displayed as a dotted line in both upper and lower panels. **(d)** The final assignment of spikes to the two putative units. Unit a is visualized in dark green; Unit b is visualized in light green. Unsorted spikes are visualized in gray.

We next examine what fraction of spikes are originating from sources other than the neuron of interest. As each spike waveform is added to a cluster, starting with those closest to the centroid and moving outward, we calculate the number of ISI violations incurred in Figure 2b. The number of ISI violations out of the total number of waveforms can then estimate the fraction of waveforms originating from the cluster of interest. Figure 2b displays that estimated fraction of true spikes which decreases as more ISI violations are encountered. As a second estimate of fraction of rogue spikes, we can also calculate overlap using our multivariate Gaussian model and *χ*^2^ fit. We use the minimum of either the ISI violations or the *χ*^2^ fit as a conservative estimate of the fraction of false positive spikes (Figure 2c upper panel).

Finally, we derived an optimization parameter, *ϕ*, for best selecting for the tradeoff between the fraction of observed spikes from the neuron of interest (*f*_*O*_) and the fraction of false spikes included (*f*_*F*_), see Methods for its derivation. Specifically, we identified that the function 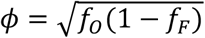 should be maximized to get the most information about a sorted unit’s firing rate. For our example channel, *f*_*O*_ and *f*_*F*_ and the resulting *ϕ* values are shown as a function of the number of spikes assigned to the units’ cluster (Figure 2c). The estimate of the best boundary of spikes to be included is indicated with a dashed line at the maximum value of *ϕ* for each sorted unit. Figure 2d plots the resulting sorting of assigned waveforms. Note that Unit c is not included in the final sort because ISI violations were encountered early near the centroid (Figure 2b, Unit c), suggesting a single neuron cannot be isolated from other neurons and noise.

### Sorting Comparison

Having shown how our spike sorting principles apply to a single channel, we next benchmarked our implementation, WaveLimit (WL), against three spike sorters: SpyKING CIRCUS (SC)^10^, IronClust (IC)^11^, and Trideclous (TC)^12^. Figure 3 captures the variegated spike assignments to the putative units found by the different sorters on the example channel from Figures 1 & 2. Waveforms that crossed each sorter’s amplitude threshold during the recordings, distinguishing them as spikes, are illustrated in Figure 3a. We assessed the unit quality by plotting inter-spike intervals (ISIs) histograms (Fig. 3b). WL unit a was congruent with IC unit a and TC unit a, while WL unit b corresponded with SC unit b and IC unit b. WL’s first unit was well isolated, exhibiting strong waveform modulation (Fig. 3a) and minimal ISI violations (no spikes near 0 in Fig. 3b). WL’s second unit posed a more challenging sorting problem, yet WL and IC performed similarly. SC identified an analogous unit, but it included significantly fewer spikes. IC and TC detected a third unit predominantly composed of ISI violations, suggesting substantial noise contamination and unlikely originating from a single neuron (this is likely the same unit as the third cluster, unit c from Fig. 2, that WL removed because of ISI violations). Thus, WL outperformed the other methods, even under difficult sorting conditions.

**Figure 3.**
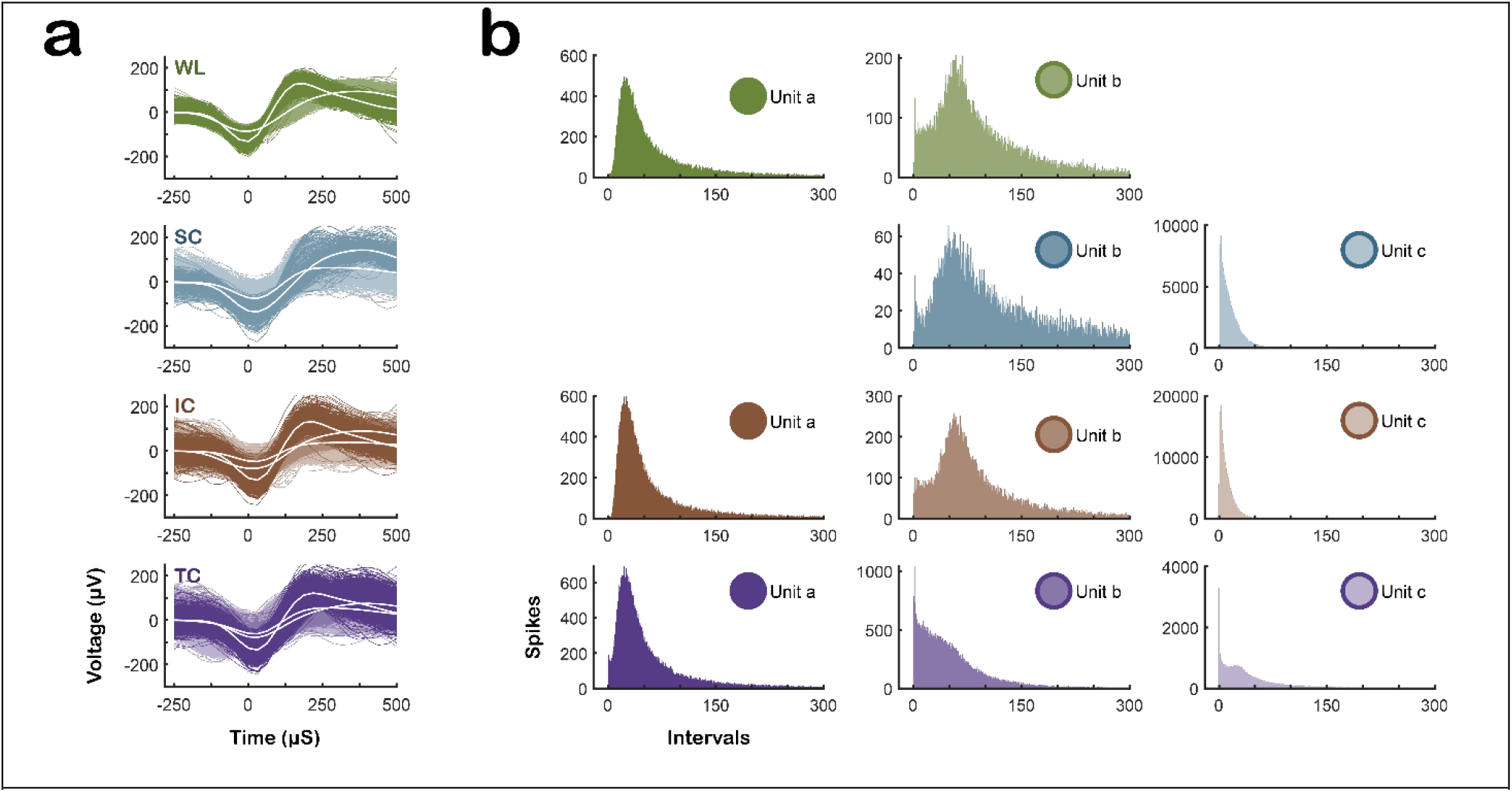
Comparison of Spike Sorting Algorithms on an Example Channel. **(a)** Waveforms of putative units identified by WaveLimit (WL), SpyKING CIRCUS (SC), Ironclust (IC), and Trideclous (TC) on an example channel. Units are colored distinctly, with the average waveform shown in white. **(c)** Inter-spike interval histogram for each unit identified. NB Only spikes assigned to units are included; unsorted waveforms are excluded.

WL identified the two units corroborated by the other algorithms’ collective sorting and excluded units dominated by ISI violations, which the others included as units. Did this balanced approach generalize well to the entire dataset, or was WL too conservative, finding fewer units and fewer spikes per unit? We counted the channels with units and tallied the total units identified by each sorter across all 768 channels, which were recorded during twelve sessions from three non-human primates each. To avoid inflation of statistical comparisons by biologically implausible units with abnormally high spike counts, we excluded extreme outliers before analysis (see Methods). After exclusion, 38 units were trimmed from SC, 70 from IC, and 7 from TC (Fig. 4a, unfilled bars). Figure 4a’s two-category bar chart reveals that WL and SC found significantly more channels with units than IC and TC (95% confidence Intervals: WL [567.23, 697.58], SC [505.65, 629.16], IC [227.71, 313.12], and TC [284.16, 378.67]). Similarly, WL and SC found more units overall (95% confidence Intervals: WL [844.71, 1002.09], SC [742.39, 890.41], IC [573.87, 704.94], and TC [456.54, 574.27]). We next examined the spike counts of the units identified, illustrated in the Raincloud plots of Figure 4b^13^. WL found more spikes per unit than SC, IC, or TC (Wilcoxon signed-rank test, *p* < 0.001, Bonferroni-corrected). Therefore, WL was not too conservative and either outperformed or rivaled the other sorters.

**Figure 4.**
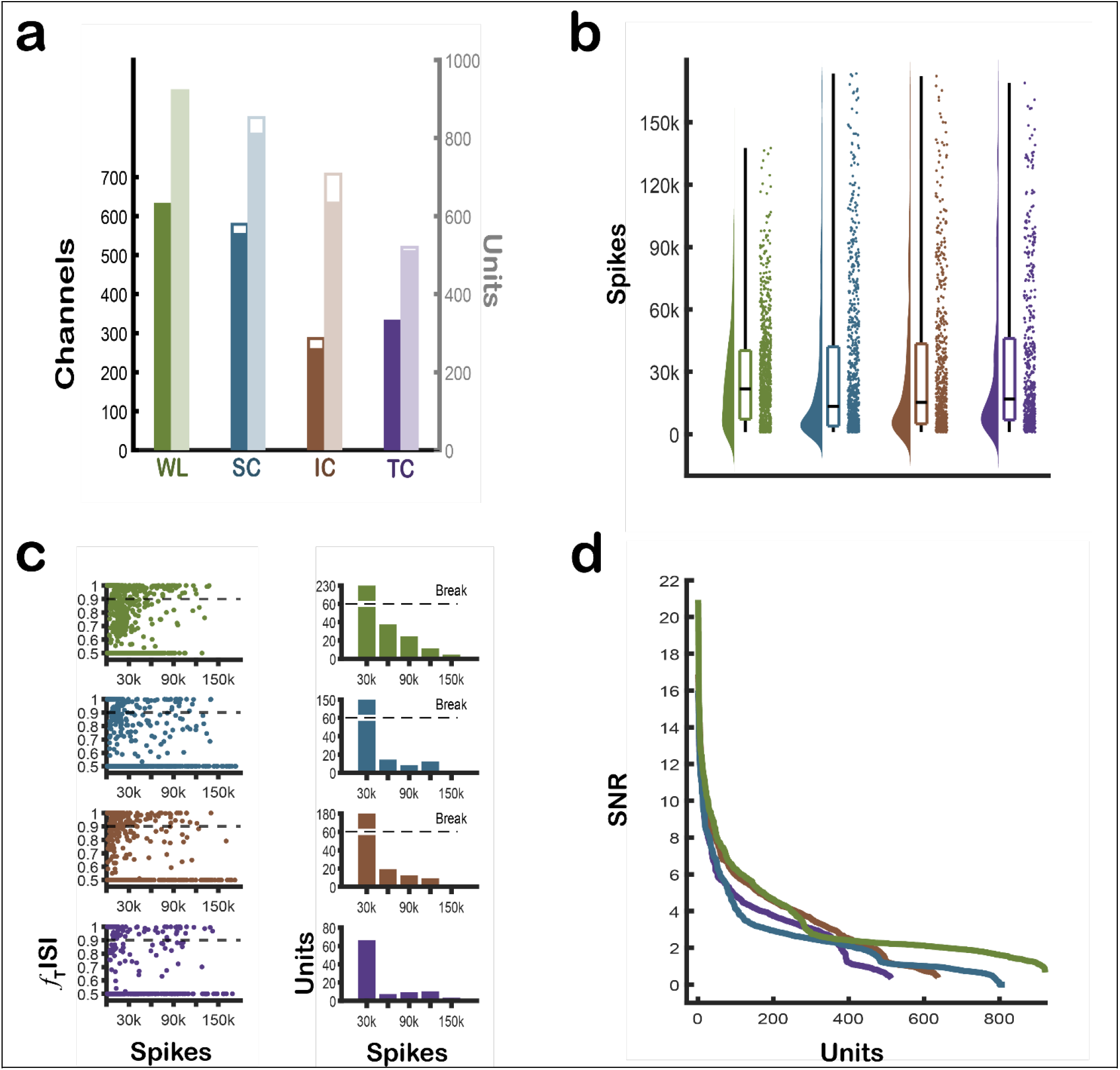
Comparison of Spike Sorting Algorithms Across All Recordings. **(a)** Darker shade represents the number of channels with at least one unit; the lighter shade represents the total number of units. Unfilled portion of each bar plot represents those units or channels that were trimmed by the outlier exclusion criteria. **(b)** Raincloud plot displays the number of spike events included in each unit. **(c)** Scatter plot displays the fraction of true inter-spike intervals (ISIs) per unit identified (true ISI ratio of 0.5 signifies no dominant single unit). Adjacent bar plots display the number of units that had a fraction of true ISIs (*f*_true_ISI) > 0.9, binned by the number of spikes included in those units. **(d)** Ranked order decay plot displays the SNR for every unit.

In Figure 3, both SC and IC identified a noisy third unit with numerous ISI violations. We therefore evaluated the integrity of the spikes included within each unit by calculating the occurrences of spikes with inter-spike intervals (ISIs) shorter than 1.5 ms^14^. The scatter plots of Figure 4c show the proportion of true spikes in each unit (identified by non-violating ISIs) as a fraction of the total number of spikes included per unit. We further considered well-isolated units to be those with less than 10% false positive spikes by number of ISI violations, shown by the horizontal dashed line on each scatter plot. We visualized the count of these well-isolated units, differentiated by the number of spikes, in the adjacent bar graphs. WL identified units with fewer ISI violations than those found by SC, IC, or TC (Wilcoxon signed-rank test, *p* < 0.001, Bonferroni-corrected)

WL was designed to maximize the spike counts in each unit while minimizing the number of ISI violations incurred. This design yields more units than the other sorters and the units identified include more spikes and fewer ISI violations. But does the design offer any additional advantages relevant for spike sorting? To explore this, we examined another feature of unit quality: the signal-to-noise ratio (SNR). We first measured the variability of each unit’s mean waveform to determine the signal component. We then subtracted the mean waveform from each individual spike waveform and calculated the standard deviation of these differences to obtain a measure of the residual noise variability. Finally, we computed the SNR as the ratio of signal variability to residual noise variability. Units identified by WL exhibited significantly higher SNRs than those of other sorters (Wilcoxon signed-rank test, *p* < 0.001, Bonferroni-corrected) (Figure 4d).

### Sorting Speed

All data was sorted on the same computer with the following specifications: an AMD Ryzen 7 3700X 8-Core Processor with a 3.6 GHz base clock, 64 GB DDR4 memory at 2133 MT/s, and an NVIDIA GeForce RTX 2070 SUPER GPU with 8 GB GDDR6. The operating system was Ubuntu 20.04.2 LTS. Table 1 outlines the processing times (raw, by unit and by total spike count) for the four spike sorters on one recording session. WL sorted the recording the fastest (892 seconds), achieved the highest rate of units per second (0.0987) and sorted the most spikes per second (4047). In contrast, SC had the longest processing time (3072 seconds). IC and TC had similar sorting rates.

**Table 1.** Sorter Processing Time. The total sort time is listed in seconds for a single recording session for each algorithm. We also list that sort time by unit and spike rates.

| Sorter | Sort Time (12 Cores) | Units / Sort Time | Spikes / Sort Time |
| --- | --- | --- | --- |
| WL | 892s | 0.0987 u/s | 4047 spikes/s |
| SC | 3072s | 0.0277 u/s | 1430 spikes/s |
| IC | 2,420s | 0.0277 u/s | 4031 spikes/s |
| TC | 2,111s | 0.0322 u/s | 1350 spikes/s |

## Discussion

We developed a novel spike sorting algorithm, WaveLimit, that automatically identifies spike waveform clusters using optimal spike inclusion boundary criteria based on established spike quality metrics. We then benchmarked WaveLimit against three sorting algorithms: SpyKING CIRCUS^10^, IronClust^11^, and Trideclous^12^. Compared to these three state-of-the-art algorithms, WaveLimit i) identified more units, ii) included more spiking events per unit with both fewer inter-spike interval violations and higher signal-to-noise ratios, and iii) sorted faster.

In recent years, spike sorting algorithms have improved progressively, especially for large channel count, high-density recordings. However, these algorithms rely on arrays that experience significant channel crosstalk – such as dense silicon CMOS probes. Many spike sorters operate by evaluating cross correlograms that are less informative in larger microwire multielectrode arrays^15^. We built WaveLimit to accommodate these larger microelectrode arrays, reducing reliance on the crosstalk expected on dense probes. Though further analysis of such recordings will be required, we anticipate WaveLimit will also perform well on other array types.

Canonically, inter-spike interval violations have been used as a *post hoc* quality metric. WaveLimit incorporates inter-spike interval violations when estimating the spike inclusion boundary. A potential limitation is that inter-spike interval violations is no longer a completely independent metric to evaluate sorting quality as WaveLimit adjusts its sorting to reduce inter-spike interval violations. However, WaveLimit demonstrate improvement in all three quality metrics tested: signal-to-noise ratios, fewer inter-spike interval violations, and spike count per unit. Although datasets can be sorted with varying emphasis on these metrics, our sorter’s improvement across all three indicates an overall sorting enhancement, not just optimization for inter-spike interval violations.

Our derivation optimizes the theoretical information available from a single neuron. Other waveform inclusion criteria may be more appropriate depending on application. If certainty of waveforms from a single neuron is required, more strict inclusion criteria will be needed. If total information regardless of neuron source is desired, more relaxed inclusion criteria might be warranted^16-18^.

Here, we concentrated on the final aspect of spike sorting: assigning an appropriate number of waveforms to a unit cluster. Thus, the principles outlined are likely not in competition with advancements made in other aspects of sorting but instead may be integrated harmoniously within much of other sorters’ existing processes. Our current version of WaveLimit employs relatively simple cluster identification and waveform distance measures^19-21^. We foresee future advances by applying our waveform inclusion techniques to other sorting frameworks or refining future versions of WaveLimit for improved unit identification and/or cluster assignment.

## Methods

### Details of the Algorithm

#### Deriving an Ideal Sorting Boundary

Choosing the proper boundary for a cluster of spike waveforms originating from a single neuron requires balancing the trade-off between missing true spikes and incorrectly including false spikes. If the boundary is too small around the cluster center, actual spikes will be excluded, underestimating the neuron’s firing rate. Conversely, if the boundary is too large, spikes from other neurons or noise will be included, overestimating a neuron’s firing rate. Both scenarios reduce the accuracy of evaluating the neuron’s response to experimental conditions and may bias the results if the missed or added spikes do not occur independently relative to the task.

Here, we derive the optimal sorting boundary for an idealized model that consists of two independent Poisson point processes. The first point process represents the observed neuron of interest with firing rate, λ. If sorting is not perfect, we can only observe a fraction of all spikes, *f*_*O*_, and the observed firing rate is *f*_*O*_λ. The second point process is an independent noise source with firing rate, λ_N_. This noise source represents all rogue spikes caused by other units or noise artifacts incorrectly classified as spikes from the neuron of interest. For a given time window, *T*, the experimentally observed spike count is the sum of all observed true spikes as well as all rogue spikes erroneously assigned to the neuron of interest:

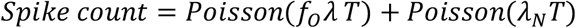

The Spike count is the sum of two Poisson distributed random variables with expected counts of *f*_*O*_*λT* from the neuron of interest and *λ*_*N*_*T* from rogue spikes.

An experimenter aims to best discriminate between different spike rates for the neuron of interest during different experimental conditions. In the simplest case, we examine discriminating the spike count during two conditions, *Spike count*_*1*_ and *Spike count*_*2*_, with different neuron firing rates, λ_1_ and λ_2_. The two conditions occur for the same length of time, *T*, and assumed to have the same noise source with a firing rate of *λ*_*N*_.

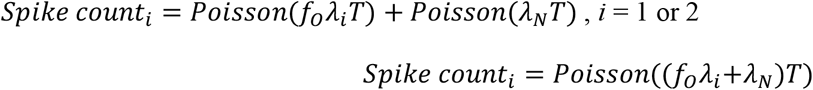

One can define the discriminability (*d’*) as the difference in means (*μ*) between the two conditions divided by the pooled standard deviation 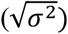:

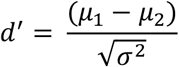

Assuming a Poisson distribution, the mean and variance are both equal to the expected firing rate. Substituting *μ*_*i*_ = (*f*_*O*_*λ*_*i*_+*λ*_*N*_)*T* and *σ*^2^ = (*f*_*O*_*λ*+*λ*_*N*_)*T*

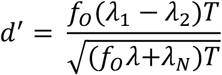

We now define the rate of rogue spikes in relation to the spikes from the neuron of interest by defining a fraction of false spikes, *f*_*F*_.

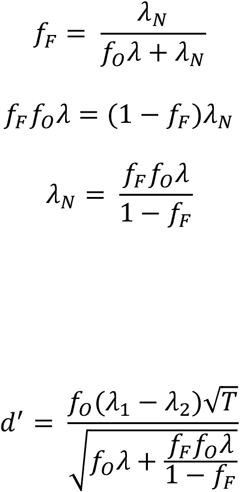

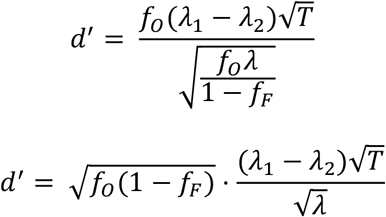

The neuron’s firing rate in the two conditions, λ_1_ and λ_2_, are intrinsic properties of the neuron and experimental conditions with discriminability increasing with increased observation time. As expected, the ability to discriminate between the two conditions also depends on the ability to maximize the fraction of spikes that are observed, f_O_, and to minimize the fraction of false spikes that are included in the observed spikes, f_F_. This result gives the appropriate trade-off between fraction of spikes observed and fraction of false spikes to maximize discrimination. Specifically, one should maximize a parameter we’ll name *ϕ*:

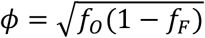

Knowing this optimal relationship, we implemented an algorithm to estimate the fraction of observed spikes and fraction of false spikes as a function of number of spikes included in a sorted single unit. Before selecting a boundary, clustering is performed to determine the units present. Our spike sorter, WaveLimit, uses the number of neighboring waveforms and a likelihood ratio test to identify cluster centers. However, numerous automated algorithms have been described and previously implemented that could also be used for cluster identification. WaveLimit also allows manual unit identification as an alternative to automatic cluster identification. For each identified cluster, either a single waveform or the mean of the labeled waveforms belonging to the cluster is defined as the characteristic waveform for each unit. Then, a distance metric is needed to rank waveforms from the waveform most likely to be a member of the cluster to waveform least likely to be included. With this ranking, it then becomes possible to choose an appropriate inclusion boundary for each sorted unit.

#### Implementation in Matlab

WaveLimit is a Matlab implementation for spike sorting using the optimal trade-off between the estimated fraction of included true spikes and fraction of false spikes using the open-source *.nex file format (NeuroNexus). The program can either automatically identify clusters or gives the user the flexibility to identify clusters either manually or using a different algorithm. The user then inputs all waveforms for a channel, the spike times of all waveforms, and the pre-identified clusters. The characteristic waveform for a cluster is calculated as the mean of all waveforms initially assigned to a given cluster.

WaveLimit then begins by calculated each waveform’s Euclidian distance from the characteristic waveforms. The Euclidian distance is calculated with:

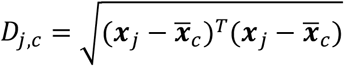

Here, ***x***_*j*_is the vector of samples over the timepoints for waveform j and 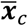 is the mean characteristic waveform.

#### Distance from mean waveform

If we assume the waveforms are created by a mean waveform with Gaussian white noise added independently to each timepoint, the squared distances from the centroid would follow a χ^2^ distribution with degrees-of-freedom equal to the number of timepoints for each waveform. In reality, the noise between adjacent timepoints in the signal is correlated which would be ideally fit with a generalized chi-squared distribution. Since there is no closed-form expression for a generalized chi-squared distribution, we found that the distribution could be well approximated with a χ^2^ distribution with reduced degrees-of-freedom and a scaling factor plus linear offset.

In our sorting algorithm, we use this χ^2^ distribution to estimate what fraction of true spikes from an individual neuron would be included for any given distance from the centroid. Since waveforms nearest the centroid are very likely to be from the neuron of interest, we aimed to estimate the 25th percentile of waveforms from the neuron of interest and then fit the χ^2^ distribution to those waveforms to estimate the entire distribution. First, we estimate the noise distribution by calculating the covariance matrix of the difference between all waveforms closer to a given characteristic waveform to estimate the degrees-of-freedom, expected mean distance, and variance. Then, using a method of moments approach, we estimate a χ^2^ distribution (with scaling and offset) that best approximates these features—degrees-of-freedom, mean, and variance—for the covariance matrix. This estimated distribution then allows us to estimate the distance from the centroid where the 25th percentile waveform would occur. Finally, using these 25 percent of waveforms, we use least-square curve fitting (lsqcurvefit in Matlab) to estimate the degrees-of-freedom, scaling, and offset of a χ^2^ distributed cumulative density function. With a χ^2^ fit established for each identified unit, the probability of a waveform originating from a given unit can be estimated. Additionally, the cumulative density function, CDF, can be used to determine what percentage of spikes would be expected to have a D value less than any distance boundary.

Our goal is to determine the optimal discrimination that can be performed by maximizing the fraction of true spikes observed, f_O_, and minimizing the fraction of false spikes, *f*_*F*_. The true spikes observed, *f*_*O*_, can be estimated by using the CDF from the χ^2^ for the observed D values.

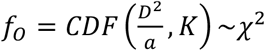

The fraction of false spikes, *f*_*F*_, is estimated from two measurements: the amount of overlap with other waveform clusters as modeled with χ^2^ distributions and the number of ISI violations. The first depends on estimating the number of spikes occurring from all waveform clusters using the CDF derived above for units other than the spike cluster of interest.

The second depended on the number of spikes with ISI violations relative to total spikes. The percent of false spikes estimated to originate from a source other than the sorted unit of interest was estimated using the number of inter-spike interval violations less than a chosen maximum refractory time (*t*_*max*_ ) in which no spikes should be generated from the same neuron.

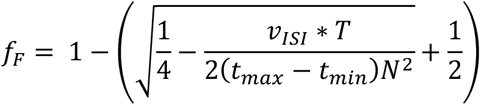

Where *v*_*ISI*_ is the number of inter-spike interval violations obverted, *N* is the total number of sorted spikes, *T* is the total recording time, *t*_*min*_ is the waveform sort width where no new spikes are detected (0.675 ms for our presented data), and *t*_*max*_ is the chosen refractory period (1.5 ms) (Hill et al., 2011). Note, whereas an absolute refractory of 1 ms for pyramidal neurons in cortex is often described, we found that increasing the chosen refractory period to 1.5 ms provided more robust performance. The spikes within this period may contain a small number of spikes during a neuron’s relative refractory period but a majority are likely contaminating rogue spikes whose observations are increased with the increased *t*_*max*_. To be conservative, we used the minimum *f*_*F*_ between the two calculations.

Once WaveLimit has calculated the *f*_*O*_and *f*_*F*_ for any distance, D, from the characteristic waveform, the D that maximizes *ϕ* is selected as the cluster boundary. All waveforms less than D are included in the final sorted unit. These sorted waveforms represent WaveLimit’s estimate for the best balance between including most of the spikes from a single neuron while not including too many rogue spikes.

Three additional features are included in the WaveLimit software package: i) Automatic cluster identification, ii) Characteristic waveform optimization, iii) Outlier rejection.

#### Automatic cluster identification

To automatically identify a cluster of waveforms that defines a presumed single unit, we first identify the center of the cluster. Our algorithm searches for waveforms that have many neighboring waveforms within a close distance to define a cluster center. We use a random sampling of 1000 waveforms per channel as candidate waveforms to be located near a cluster center. The first 500 are sampled entirely at random. Two successive groups of 250 waveforms are then added that are not near any of the original test waveforms. This successive approach helps identify small clusters of waveforms that might not be found if one large cluster has many more waveforms than a smaller cluster but remains computationally fast with only 1000 waveforms per channel compared to considering all waveforms.

We then rank the candidate central waveforms by number of close neighboring waveforms. We chose to use a likelihood-ratio test for this ranking. Using the first time sample (before future threshold crossings), we can estimate the standard deviation, σ, of noise in the neurophysiology recording system. This provides an estimate of the expected multivariate Gaussian noise distribution around the central waveform shape. We then used a likelihood ratio test to see which waveforms appear to be at the center of a cluster with Gaussian noise with standard deviation, σ. We compared this to a broader distribution of spikes around a central waveform, 2.5 σ.

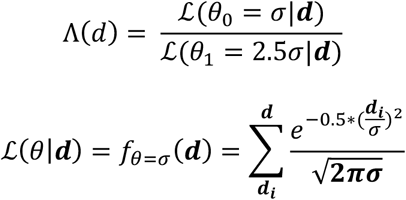

Here, **d** is a vector of all distances between all other waveforms and the candidate central waveform. The likelihood function utilizes the multivariate normal probability distribution function calculated for the distances from the characteristic waveform and tested for two different standard deviations, σ or 2.5 σ. Using the ratio test helps identify waveform clusters with the expected narrow distribution of a neuron’s same action potential waveshape occurring repeatedly plus the observed noise for each recorded individual spike. The ratio test helps to normalize for different number of waveforms in various clusters.

The candidate waveform with the highest ratio is deemed most likely to lie at the center of a cluster with multivariate Gaussian noise equal to σ. This candidate waveform is selected as the characteristic waveform to define the cluster center and any other candidate waveforms that have significant overlap with an already selected cluster center are eliminated. This iterative process is continued until all candidate waveforms are either selected as a cluster center or eliminated. A final check is performed for potential overlap between cluster center waveforms and any with too much overlap with other waveforms are eliminated.

#### Characteristic waveform optimization

A characteristic waveform optimization algorithm is included which utilizes ISI violations to adjust the characteristic waveform relative to the initial, either user supplied or identified with automatic clustering. This optimization presumably moves the characteristic waveform away from other contaminating units or noise. This iterative algorithm utilizes mean waveshape of the first few (default = 10) waveforms with observed ISI violations to shift the characteristic waveform away from these waveforms causing ISI violations. Note, since an ISI violation is comprised of two rapidly occurring waveforms and presumably one is a true spike and the other the violating spike, the 2^nd^ waveform added to the unit’s sorting—the one the farther distance from the characteristic waveform—is assumed to cause the ISI violation and treated as the ISI violating waveform. The default shift with each iteration is 10% of the distance between the mean ISI violating waveforms and the previous characteristic waveshape in the opposite direction away from the violating waveforms. The sorter iteratively adjusts the characteristic waveform until the sorting parameter, *ϕ*, no longer improves.

#### Outlier rejection

Outlier rejection based on the non-linear energy of the waveforms. The non-linear energy measures the amount of sharp discontinuities in each individual waveform. An increased non-linear energy is often a sensitive identifier of waveforms that appear less physiologic and likely caused by electrical noise^22^. WaveLimit calculates this non-linear energy for all waveforms assigned to a sorted unit and throws out all waveforms greater than a number of standard deviations (default = 5σ) from the sorted unit population.

### Data

#### Subjects

Three male rhesus macaques were subjects in the present study: P, Q, and A (weights: 11, 10, and 12 kg; ages: seven, six, and six years old, respectively). All procedures for the care and use of these non-human primates followed the *Guide for the Care and Use of Laboratory Animals* and were approved by the University Committee on Animal Resources at the University of Rochester Medical Center, Rochester, NY, (for subjects P and Q) and by the Institutional Animal Care and Use Committee at the University of Kansas Medical Center, Kansas City, KS (for subject A).

#### Neural Recordings

Floating microelectrode arrays (MicroProbes for Life Science) were implanted to record from the primary motor cortex (M1) in subjects P and Q, using methods described in detail previously^23-25^. For subject P, recordings were collected from four 16-channel arrays implanted in M1. For subject Q, two 32-channel arrays in M1 were used. A similar procedure was performed in subject A using floating microelectrode arrays (MicroProbes for Life Science). Subject A’s recordings were collected from four 32-channel arrays implanted in M1 in the anterior bank of the central sulcus. For subjects P and Q, 30-kHz broadband neural signals were collected with Cerebus (Blackrock Microsystems, LLC.) data acquisition system. For subject A, 30-kHz broadband neural signals were collected with a SpikeGadgets data acquisition system.

A precision center-out task was performed by subjects P and Q as described in detail previously^25^. A similar precision center-out task was performed by subject A using a non-human primate exoskeleton robot (KINARM, BKIN Technologies, Kingston, ON, Canada) to capture kinematic data. However, the task performed by subject A involved perturbations in which the targets toward which the subject reached were moved in a fraction of trials. Each subject performed their respective task during three sessions, recorded over a month period.

### Other Spike Sorting Algorithms

To compare WaveLimit against existing methods, we selected state-of-the-art algorithms based on the results of the SpikeForest project^26^, which identified top-performing spike sorting algorithms across standardized benchmarks. From this project, we chose SpyKING CIRCUS^10^, IronClust^11^, and Trideclous^12^ due to their high performance and lack of significant implementation issues. All algorithms processed the same recordings as WaveLimit, using the SpikeInterface^27^ platform to ensure that each spike sorter was run in the same way. All parameters were set to their default values.

### Statistical Analysis

To exclude extreme outlier units with biologically implausible spike counts (e.g., >300,000 spikes), we applied a robust nonparametric method. Specifically, we pooled all units identified by all algorithms and calculated the median and Median Absolute Deviation (MAD)^28^ of spike counts across this combined dataset. Units with spike counts exceeding the pooled median plus ten times the MAD were then removed from each algorithm’s dataset. All subsequent statistics were computed from the units that survived this outlier exclusion criterion.

95% confidence intervals for the number of channels with units and the total number of units were calculated assuming a Poisson distribution using the chi-squared method, where:

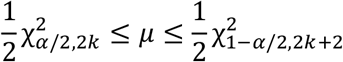

Here, *k* is the observed count, and 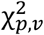 denotes the chi-squared quantile function with *p* probability and *ν* degrees of freedom. All confidence intervals were calculated at the 95% level (*α = 0*.*05*).

We used Wilcoxon signed-rank tests to compare the number of spikes per unit, the fraction of true spikes (ISIs > 1.5 ms), and the Signal-to-noise ratios (SNRs) between WaveLimit and each of the other spike sorters individually. This nonparametric test was chosen due to the non-normal distribution of the data. Because these involved multiple pairwise comparisons, Bonferroni correction was applied, with the significance threshold adjusted based on the number of comparisons performed. Unless otherwise stated, p-values are reported after correction. Statistical significance was set at *p* < *0.05* after correction.

## Software Availability

MATLAB code for WaveLimit is available publicly on GitHub^9^. Additional analysis code is available from the authors on request.

